# Distinct mechanisms drive plant-nitrifier interactions in topsoil and subsoil

**DOI:** 10.1101/2023.12.01.569677

**Authors:** Di Liang, Niuniu Ji, Angela Kent, Wendy H. Yang

## Abstract

Plants can influence soil microbes through resource acquisition and interference competition, with consequences for ecosystem function such as nitrification. However, how plants alter soil conditions to influence nitrifiers and nitrification rates remains poorly understood, especially in the subsoil. Here, coupling the ^15^N isotopic pool dilution technique, high throughput sequencing and *in situ* soil O_2_ monitoring, we investigated how a deep-rooted perennial grass, miscanthus, versus an adjacent shallow-rooted turfgrass reference shapes nitrifier assembly and function along 1 meter soil profiles. In topsoil, the suppression of ammonia (NH_3_) oxidizing archaea (AOA) and gross nitrification rates in miscanthus relative to the reference likely resulted from nitrifiers being outcompeted by plant roots and heterotrophic bacteria for ammonium (NH_4_^+^). The stronger tripartite competition under miscanthus may have been caused in part by the lower soil organic matter (SOM) content, which supported lower gross nitrogen (N) mineralization, the major soil process that produces NH_4_^+^. In contrast, below 10 cm soil depth, significantly greater gross nitrification rates were observed in miscanthus compared to the reference. This was likely driven by the significantly lower oxygen (O_2_) in miscanthus than reference subsoil, which selected against aerobic heterotrophic bacteria but in favor of AOA. Overall, we found that plants can regulate AOA community structure and function through different mechanisms in topsoil and subsoil, with suppression of nitrification in topsoil and enhancement of nitrification in subsoil.

## 1. Introduction

Nitrogen (N) is a major limiting nutrient for plant growth in most ecosystems (Tilman, 1984). To control soil N retention in the plant-soil interface, plants have developed multiple strategies such as resource acquisition and interference competition to regulate N-cycling microbes, with consequences for ecosystem function such as nitrification (Kaye and Hart, 1997; Moreau et al., 2019). Nitrification plays a central role in the global N cycle by regulating the availability of N to microbes and plants, and also increasing the potential for negative environmental impacts such as nitrate (NO_3_^-^) leaching and emissions of nitrous oxide (N_2_O), a potent greenhouse gas with a global warming potential 300 times higher than carbon dioxide (CO_2_) (Prosser, 1990; Robertson and Groffman, 2015). Mediated mainly by ammonia (NH_3_) oxidizing archaea (AOA) and bacteria (AOB) and complete NH_3_ oxidizers (comammox) (Prosser and Nicol, 2012; Daims et al., 2015; van Kessel et al., 2015), nitrification is a microbial process that converts NH_3_ to nitrite (NO_2_^-^) and NO_3_^-^. Through exploitative competition, plant roots can decrease soil nitrification by reducing the quantity of NH_4_^+^ available for nitrifiers (Zak et al., 1990; Hawkes et al., 2007). Moreover, some plant species, such as tropical grass *Brachiaria*, can release root exudates that directly inhibit nitrification (Subbarao et al., 2009).

Despite the intensive efforts in understanding plant-nitrifier interactions, current research has mostly focused on topsoil, where root biomass is greatest (Gill et al., 1999). However, plant-microbe interactions in subsoil can potentially be as intense as topsoil, particularly under deep-rooted plants (Jones et al., 2018). Whether the environmental and plant drivers of soil microbial processes, such as nitrification, are similar or different between topsoil and subsoil remains uncertain.

Physical and chemical properties of subsoil often significantly differ from those of topsoil, resulting in distinct regulatory mechanisms for biogeochemical processes and microbial communities through soil profiles (Salomé et al., 2010; Tess et al., 2019). A recent deep metagenomic sequencing and metaproteomic analyses demonstrated that both microbial phylogenetics and functionality were highly stratified with soil depths (Diamond et al., 2019). While microorganisms encoding carbon metabolisms and degradation were enriched in surface soil, microbial N turnover pathway such as NH_3_ oxidation was more prevalent in deep soil. The more abundant C turnover functions seem to reflect the accumulation of organic matter derived from plant litter and root exudates in upper profiles (Lennon, 2020). Meanwhile, in deeper horizons that have much less plant inputs of soil organic carbon and N (Button et al., 2022), the increased abundance and functions of AOA have been frequently attributed to their abilities to thrive in oligotrophic conditions. This differs from AOB as the growth of AOB is typically favored in fertilized systems (Ouyang et al., 2016; Liang et al., 2020) or in the nutrient-enriched surface soil (Banning et al., 2015; Mushinski et al., 2017).

Aside from the niche separation between AOA and AOB (Prosser et al., 2019; Trivedi et al., 2019), the enrichment of AOA in subsoil could also be derived from the competition between heterotrophic bacteria and nitrifiers for available NH_4_^+^ and oxygen (O_2_) (Nogueira et al., 2002). Due to the low substrate requirement and the insensitivity to O_2_ limitation (Pett-Ridge et al., 2013; Ke et al., 2015; Wang et al., 2015), AOA outcompeted heterotrophic bacteria and dominated the microbial community in the hypoxic zone (18 m below sea level) in the Gulf of Mexico (Gillies et al., 2015). However, the O_2_ mediation of AOA-heterotroph competition has not been explored in terrestrial ecosystems, although rhizosphere priming of microbial O_2_ consumption could deplete soil O_2_ in the subsoil where atmospheric O_2_ exchange is slow (Philippot et al., 2009; Lennon, 2020), thereby favoring AOA and potentially increasing nitrification rates. This presents the potential for plants to influence soil nitrifier community structure and function via different mechanisms in topsoil versus subsoil.

To better understand how plants alter soil conditions to influence nitrification in topsoil and subsoil, we investigated the function and community structure of soil bacteria and nitrifiers down 1 m soil depth profiles in mature (10-year-old), deep-rooted miscanthus (*Miscanthus × giganteus*) stands (miscanthus soil) and adjacent shallow-rooted turfgrass references (reference soil). Because of the differential nutrient uptake and the various effects of plants on environmental variables along soil profiles, we hypothesized that deep-rooted plants would employ distinct mechanisms to regulate nitrifiers and their surrounding microbes in topsoil vs. subsoil. To test this hypothesis, we combined the ^15^N isotopic pool dilution technique, high throughput sequencing and measurement of soil physiochemical properties known to affect nitrification, including *in situ* soil O_2_ concentrations (Fig. S1), along the soil profiles.

## 2. Materials and methods

### 2.1 Site description

This study was conducted at the University of Illinois Energy Farm in Urbana, Illinois (40°04′ N, 88°13′ W), on a foot slope position in the Embarras River Watershed. The site has Drummer silty clay loam soils (fine-silty, mixed, superactive, mesic Typic Endoaquolls), with land capability class (LCC) of 5W (USDA NRCS, 2001) due to frequent spring flooding that limits normal agricultural productivity (Zumpf et al., 2019). The mean annual temperature and precipitation between 2010-2019 for the area were 9.0 ℃ and 1023 mm, respectively (Burnham et al., 2022).

A large bioenergy feedstock trial comparing different perennial grass feedstocks, harvesting times, and fertilization rates was established at the study site in 2013. The experiment is arranged as a split-plot within a randomized complete block design with four replicates, including three harvesting times as the large plot factor, five different perennial grass feedstocks within each harvest regime, and three nitrogen fertilization rates within each feedstock as the subplot and sub-subplot factors (Zumpf et al., 2019). To characterize the effects of deep-rooted plants in this study, we used the unfertilized miscanthus (*Miscanthus x giganteus*) sub-subplots (1.5 m x 4.6 m) harvested after frost killing, which has been shown to be the preferred harvest practice for miscanthus as it reduces nutrient removal while optimizing biomass yield potential (Cooney et al., 2022). The miscanthus was planted by plug with 60 cm row spacing. We used the unmanaged turfgrass alleyways behind the large plots to characterize the effects of shallow-rooted plants as a reference comparison to elucidate miscanthus effects in the subsoil.

### 2.2 Soil sampling

Soil samples were collected in May 2021 to characterize the rhizosphere soil microbiome structure and function when plants were active in both the miscanthus plots and the reference alleyways. We removed litter before sampling soil down to 1 m depth using hydraulic soil coring probes (Giddings Machine Co., Windsor, CO, USA) mounted to small tractors (Model 5065E with 512 front loader, John Deere, Moline, IL, USA). Three sub-replicate soil cores (5.7 cm outer diameter) were randomly collected from each miscanthus plot and from the reference alleyway adjacent to each plot (n = 4). The soil cores were divided into five depth intervals in the field: 0-10, 10-20, 20-30, 30-50 and 50-100 cm. The three sub-replicates of each depth interval were composited and stored in a cooler with ice. Immediately after transporting soil back to the lab, soil was divided into subsamples: unsieved soil was used for measuring gross microbial process rates at room temperature (25 °C); 2 mm sieved soil stored at 4 ℃ was for measuring soil properties; and 2 mm sieved soil stored at -20 ℃ freezer was used for DNA-based characterization of the microbial community. Throughout the paper, we defined topsoil as 0-10 cm depth and subsoil as 10-100 cm soil depth (Button et al., 2022).

### 2.3 Soil microbiome characterization

To characterize the soil microbiome structure and function, soil DNA was extracted from 0.25 g freeze-dried soil using the DNeasy PowerSoil Pro Kit (QIAGEN, Germantown, MD, USA). DNA concentration was determined by Qubit 4 Fluorometer (Invitrogen, Carlsdab, CA, USA). DNA quality was analyzed using NanoDrop One Microvolume UV-Vis Spectrophotometer (Thermo Scientific, MA, USA). Amplicon sequencing of V4 region of 16S rRNA, *amoA* gene of AOA and AOB were conducted in the Roy J. Carver Biotechnology Center at the University of Illinois at Urbana-Champaign. We used a 2-step PCR protocol recommended by Fluidigm Access Array system (Fluidigm, San Francisco, CA) to prepare DNA amplicons at an annealing temperature of 55 ℃. The initial PCR was implemented to amplify DNA region using primers of 515F/806R for 16S rRNA (Apprill et al., 2015; Parada et al., 2016), *amoA-*1F/*amoA*-2R for AOB (Rotthauwe et al., 1997) and *CrenamoA*-23F/*CrenamoA*-616R for AOA (Tourna et al., 2008), with Fluidigm-specific primer pads of CS1 (5′- ACACTGACGACATGGTTCTACA -3′) and CS2 (5′-TACGGTAGCAGAGACTTGGTCT-3′).

In the second step, 1 µL of 100-fold diluted product from the initial PCR was used as a template. In this step, Illumina adaptors of i5 (5′-AATGATACGGCGACCACCGAGATCT-3′) and barcoded i7 (5′-CAAGCAGAAGACGGCATACGAGAT-3′) were incorporated. Final amplicons were quantified, purified, and combined to the same concentration before sequencing with Illumina NovaSeq 6000 with 2 x 250 bp paired-end chemistry.

To process 16S rRNA sequences, we first trimmed primers from the demultiplexed raw sequences using FASTX-Toolkit (Hannonlab, 2010). Then paired-end sequences were merged. Merged reads with length < 230 bp, and sequences containing ambiguous bases “N” were discarded. We further filtered out low quality sequences that had quality scores lower than 30 in more than 10% of the bases. VSEARCH (v. 2.4.3) (Rognes et al., 2016) was subsequently used to dereplicate filtered sequences, remove singletons, and cluster operational taxonomic units (OTU) at 97% similarity. Chimeras were first removed *de novo*, followed by referencing against the “Gold” database (http://sourceforge.net/projects/microbiomeutil/files/, accessed May 1, 2022). Finally, OTU table was generated with VSEARCH, and taxonomy was assigned via the REference Sequence annotation and CuRatIon Pipeline (RESCRIPt, (Pertea et al., 2021)) prepared Naïve Bayes classifier for the V4 region (Bokulich et al., 2018) against the SILVA database (v. 138.1) (Quast et al., 2013). Once chloroplast and mitochondrial OTUs were removed, read counts were eventually rarefied to 71023 to correct for differential sequencing depth (Fig. S2). Relative abundance of AOA was based on the frequency of class *Nitrososphaeria* of phylum *Thaumarchaeota* in 16S rRNA genes excluding Group 1.1C-associated sequences. Relative abundance of AOB was based on the frequency of genera *Nitrosomonas* sp. and *Nitrosospira* sp. in 16S rRNA genes.

For *amoA* AOA and AOB, raw sequences were similarly processed as 16S rRNA sequences except taxonomy assignment. High quality *amoA* reference sequences with minimum score of 463 and size larger than 210 amino acids for AOA, and minimum score of 333 and size larger than 160 amino acids for AOB were first downloaded from FunGene database (Fish et al., 2013). Representative OTUs of AOA and AOB were then BLASTed against dereplicated reference sequences, and OTUs with E-value greater than 1×10^-50^ and minimum query coverage below 50 were deleted. The rest of the representative OTUs were manually BLASTed against NCBI database to confirm accuracy. The taxonomy of AOA was annotated based on a global *amoA* AOA reference database collected by Alves et al. (2018). Read counts of AOA and AOB were eventually rarefied to 1455 and 1916 to correct for differential sequencing depth (Fig. S3, S4).

### 2.4 Microbial process rate quantification

To directly assess function of the ammonia-oxidizing communities, we measured gross nitrification rates using the ^15^N stable isotope pool dilution technique (Krichels et al., 2019). We also used the same technique to measure gross N mineralization rates to estimate N supply from soil organic matter. Briefly, about 100 g of unsieved soil was placed in a Ziploc bag. Then, either 1 ml of 99 atom% K^15^NO_3_ containing 11.7 µg N ml^-1^ or 1 ml of 99 atom% ^15^NH_4_Cl containing 11.0 µg N ml^-1^ was evenly added to the soil using a pipette and gently mixed to evenly distribute the ^15^N label in the soil. The ^15^N label solution concentrations were used to target 10 atom % ^15^N enrichment of the soil NO_3_^-^ and NH_4_^+^ pools based on pre-experiment measurements of soil NH_4_^+^ and NO_3_-concentrations. Thirty minutes after ^15^N label addition, 30 g of soil was extracted with 120 ml of 2M KCl to determine the initial concentration and ^15^N enrichment of the soil NO_3_^-^ and NH_4_^+^ pools. The remaining soil was sealed in a 473 ml Mason jar. After 24 hours, 30 g of soil in the jar was extracted with 120 ml of 2M KCl to determine the final concentration and ^15^N enrichment of the soil NO_3_^-^ and NH_4_^+^ pools.

Soil NH_4_^+^ and NO_3_^-^ concentrations were determined from colorimetric analysis of the KCl extracts using a SmartChem 200 discrete analyzer (Unity Scientific, MA, USA). ^15^N enrichment of the soil NH_4_^+^ and NO_3_^-^ pools was determined from acid trap diffusion of the KCl extracts (Herman et al., 1995), and analysis of the filter disks on an IsoPrime 100 isotope ratio mass spectrometer (IRMS) interfaced to a Vario Micro Cube elemental analyzer (Isoprime Ltd., Cheadle Hulme, UK; Elementar, Hanau, Germany). Gross mineralization and gross nitrification rates normalized to dry soil mass were calculated from the change in ^15^NH_4_ ^+^ and ^15^NO_3_ ^-^ pool sizes, respectively, using the isotope pool dilution equations as described by Hart et al. (1994).

### 2.5 Soil property characterization

We measured a suite of soil physical and chemical properties that may affect the soil microbial community structure and function (Table S1). Soil pH was measured in soil slurries at 1:2 ratios of fresh soil mass to deionized water volume using a pH meter (Orion Star A211, Thermo Scientific, MA, USA). Soil moisture was measured by oven drying 15 g of soil at 105 ℃ for 24 hours until constant weight. Soil texture was measured using the hydrometer method with 5% sodium hexametaphosphate (Gee and Bauder, 1986). Soil bulk density was measured by collecting additional deep soil cores using hydraulic probes. Soil cores were first divided into five depth intervals as previously described, followed by removing gravel and large plant debris. Soil was then oven-dried for 48 hours at 60 ℃ degree until constant weight. The gravel-free bulk density was calculated as the dry mass of soil divided by the volume of the cores. Soil water-filled pore space (WFPS) was calculated from soil moisture and soil bulk density, assuming soil particle density of 2.65. Soil organic matter content was measured using the loss on ignition method at ignition conditions of 550 ℃ for 3 hours (Hoogsteen et al., 2015). Soil NH_4_^+^ and NO_3_^-^ were extracted with 2M KCl and measured colorimetrically using the SmartChem 200 discrete analyzer as previously described.

We also measured *in situ* bulk soil oxygen concentrations along the 1 m depth profile in each miscanthus plot and paired turfgrass reference using soil gas equilibration chambers (Fig. S1). The equilibration chambers consisted of 50 mL centrifuge tubes attached to corrosion-resistant aluminum tubing (3.2 mm outside diameter) with compression fittings and rubber septa sealing the other end of the tubing, a modification of the design from Silver et al. (1999). In early December 2021, the equilibration chambers were installed vertically at 10, 20, 30, 50 and 100 cm depth in the field, with open end of the centrifuge tube at the specified depths. Gas samples were collected bi-weekly from December 2021 to April 2022, with 10 mL gas samples stored in N_2_-flushed crimp top glass vials. Soil O_2_ concentrations in the gas samples was measured using a thermal conductivity detector on a gas chromatograph equipped with a 5A molecular sieve column for separation of N_2_, O_2_, and argon (Shimadzu GC-2014, Shimadzu Scientific Instruments, Inc, Kyoto, Japan).

### 2.6 Statistical analysis

All the statistical analysis was performed in R (version 4.1.3). To investigate how AOA and AOB communities differ between miscanthus and reference soil, principal coordinate analysis (PCoA) was conducted based on the log-chord distance of the normalized OTU data using vegan package (Borcard et al., 2018; Oksanen et al., 2022). Permutational multivariate analysis of variance (PERMANOVA) was performed to test for compositional differences between miscanthus and reference soil or among different soil depths. Since no significant differences for AOB communities were detected between plant types and depth intervals, further analysis focused on the assembly and function of AOA. To analyze the effects of miscanthus on gross nitrification, a statistical model was constructed including two plant types × five soil depths, with the interactions considered a fixed factor. Field replicates nested within plant types were considered a random factor. Analysis of variance (ANOVA) was implemented using lme4 package (Bates et al., 2015) by considering plant types as a whole plot factor and soil depths as a subplot factor. We used lsmeans package to compare nitrification rates between miscanthus and reference for each soil depth as well as compare nitrification rates among depths within each soil profile (Lenth, 2016). Normality of residuals was inspected by plotting residuals against fitted values, with no violations of assumptions detected. Statistical models for gross mineralization and soil properties were constructed similarly.

To determine the relationship between soil properties and AOA community assembly, we first calculated the Bray-Curtis dissimilarity of AOA between miscanthus and reference soil for each depth. We then calculated the depth-wise Euclidean distance of each soil property variable between miscanthus and reference, after scaling and centering each variable. A random forest model was constructed by considering the Bray-Curtis distance of AOA as the response vector, and the Euclidean distance of soil property variables as predictors using the randomForest package (Breiman, 2001). The relative importance of each variable in shaping the AOA differences between miscanthus and reference was reported as an increase in node purity, with more important variables having higher increases in node purities.

To examine the interactions between AOA and heterotrophic bacteria in subsoil, we first predicted the function of prokaryotic OTUs using Functional Annotation of Prokaryotic Taxa (FAPROTAX) (Louca et al., 2016). FAPROTAX predicts function based on literature of cultured representatives, and thus results reflect the cumulative abundance of OTUs assigned to each biogeochemical pathway. The functional abundance of nitrification (constrained to AOA) and aerobic heterotrophy were calculated by dividing the abundance of each function in a sample to the total abundance of all the function in that sample. Functional abundance of AOA-derived nitrification was regressed against the functional abundance of aerobic heterotrophy using the lm function in R. The differences in regression slopes between miscanthus and reference soil were tested using analysis of covariance (ANCOVA) with lstrends function in R.

To identify the differentially abundant AOA and aerobic heterotrophic bacteria between miscanthus and reference subsoil, we performed a differential abundance (DA) test on AOA *amoA* and aerobic chemoheterotroph OTUs using the DESeq2 package (Love et al., 2014). We also applied a zero-inflated negative binomial (ZINB) model to account for the excess of zeros, typical of microbiome data. Zero inflation weights were first calculated to reduce the overestimation of dispersion hence the loss of statistical power (Van den Berge et al., 2018). We then implemented a likelihood ratio test (LRT) using paramedic fit after estimation of size factors. OTUs that were significantly enriched or depleted in miscanthus than reference soil were identified according to log2-fold changes.

## 3. Results

### 3.1 Nitrifier community structure and function

Our PCoA analysis revealed that AOA communities changed with both soil depth and plant type (*P* < 0.001). Under miscanthus, AOA community structure did not differ among the three depth intervals in the top 30 cm of soil (Fig. 1a). In contrast, OTU clusters of AOA for the reference soil varied along the 1 m soil profiles (*P* < 0.001), with the exception of the intermediate depth of 30-50 cm, which had communities similar to the adjacent depths of 20-30 cm and 50-100 cm (Fig. 1b). Additionally, the relative abundance of AOA in miscanthus was significantly lower than that of reference soil in 0-10 cm but not in other depths (*P* < 0.05, Fig. S5). For AOB, low read counts (< 100) were found below 30 cm depth for both miscanthus and reference soil, consistent with the depth-pattern of relative abundance of AOB (Fig. S5). In addition, the AOB communities were similar between both plant type and across all soil depths (0-30 cm; Fig. 1c, 1d).

**Fig. 1.**
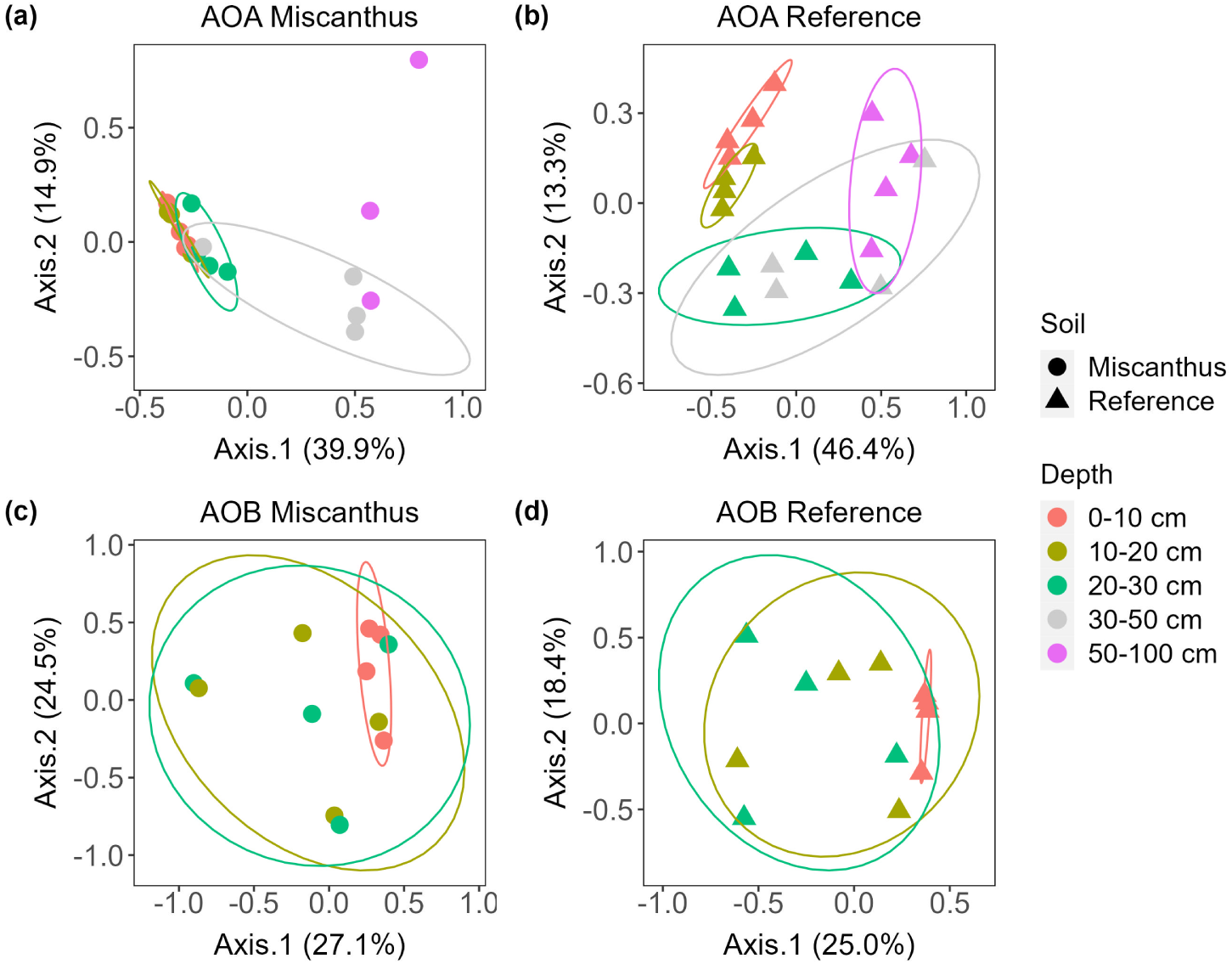
Principal coordinate analysis (PCoA) based on the log-chord distance of AOA and AOB for miscanthus (circle) and reference (triangle) soil along the 1 m profiles. Colors indicate soil depths. Ellipses correspond to 80% normal confidence interval for each soil depth interval. The proportion of variance explained by each principal coordinate is indicated on the axes’ labels. Read counts were low (< 100) for AOB below 30 cm depth so these were excluded from the analyses.

Gross nitrification rates showed distinct patterns along the 1 m soil profiles for miscanthus versus reference (Fig. 2). For the miscanthus soil, gross nitrification rates peaked at an intermediate depth of 20-30 cm, reaching a mean (± standard error) of 0.38 ± 0.19 mg N kg^-1^ day^-1^. This was an order of magnitude higher than the rates at 0-10 and 50-100 cm depths (*P* < 0.01), which averaged 0.03 and 0.02 mg N kg^-1^ day^-1^, respectively. For the reference soil, gross nitrification rates generally declined along the soil profile, with the highest rates of 0.30 ± 0.08 mg N kg^-1^ day^-1^ observed in the topsoil (0-10 cm) and the lowest rates of 0.06 mg N kg^-1^ day^-1^ observed at 50-100 cm depth (*P* < 0.05). Miscanthus had significantly lower gross nitrification rates than the reference soil at 0-10 cm depth, but this pattern was reversed below 10 cm, with gross nitrification rates of miscanthus triple that of the reference soil at the 20-30 cm depth interval (*P* < 0.05, Fig. 2). Gross nitrification rates were significantly correlated with both AOA richness and AOA relative abundance (ρ = 0.41, *P* < 0.05, Fig. S6). In contrast, no correlations between gross nitrification and relative abundance of AOB or AOB richness were observed.

**Fig. 2.**
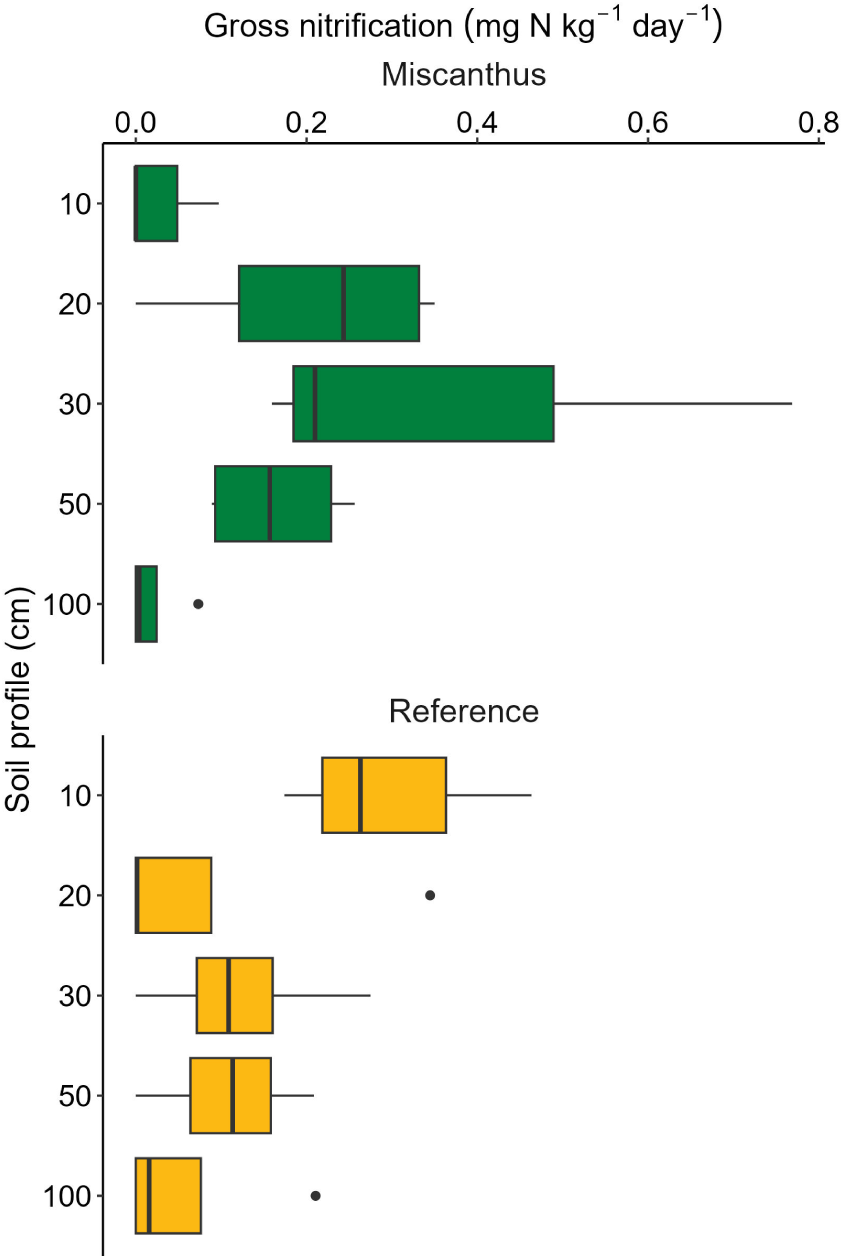
Gross nitrification rates for miscanthus (upper, green) and reference soil (lower, yellow) along 1 m soil depth profiles. The bars refer to rates measured in 0-10, 10-20, 20-30, 30-50, and 50-100 cm depth intervals. The upper, mid, and lower lines of each boxplot indicate 25^th^, median, and 75^th^ percentiles. The upper and lower whiskers indicate 1.5 × inter-quartile range (IQR), and dots indicate outliers.

### 3.2 Soil and microbial factors regulating AOA communities

The random forest model demonstrated that, among all the soil variables considered, SOM and soil O_2_ were the two most important factors in shaping the AOA community differences between miscanthus and reference soil (Fig. 3a). SOM content was lower under the deep-rooted miscanthus than the shallow-rooted reference across the 1 m soil profiles, with the greatest difference at 0-10 cm depth (8.2% vs. 10.6%, *P* < 0.01, Fig. 3c). Gross N mineralization rates demonstrated similar patterns across the 1 m soil profiles, with the greatest difference at 0-10 cm depth (6.58 vs. 10.30 mg N kg^-1^ day^-1^, *P* < 0.01, Fig. 3d). No significant plant type differences in gross N mineralization rates were detected within specific depth intervals below 10 cm. Along the 1 m soil profiles, gross N mineralization rates were significantly correlated with SOM content (ρ = 0.73, *P* < 0.001, Fig. S7). Soil O_2_ concentrations diverged between plant types only at intermediate soil depths (Fig. 3b), with lower soil O_2_ under miscanthus than reference at depth intervals of 20-30 cm (*P* = 0.01) and 30-50 cm (*P* = 0.07).

**Fig. 3.**
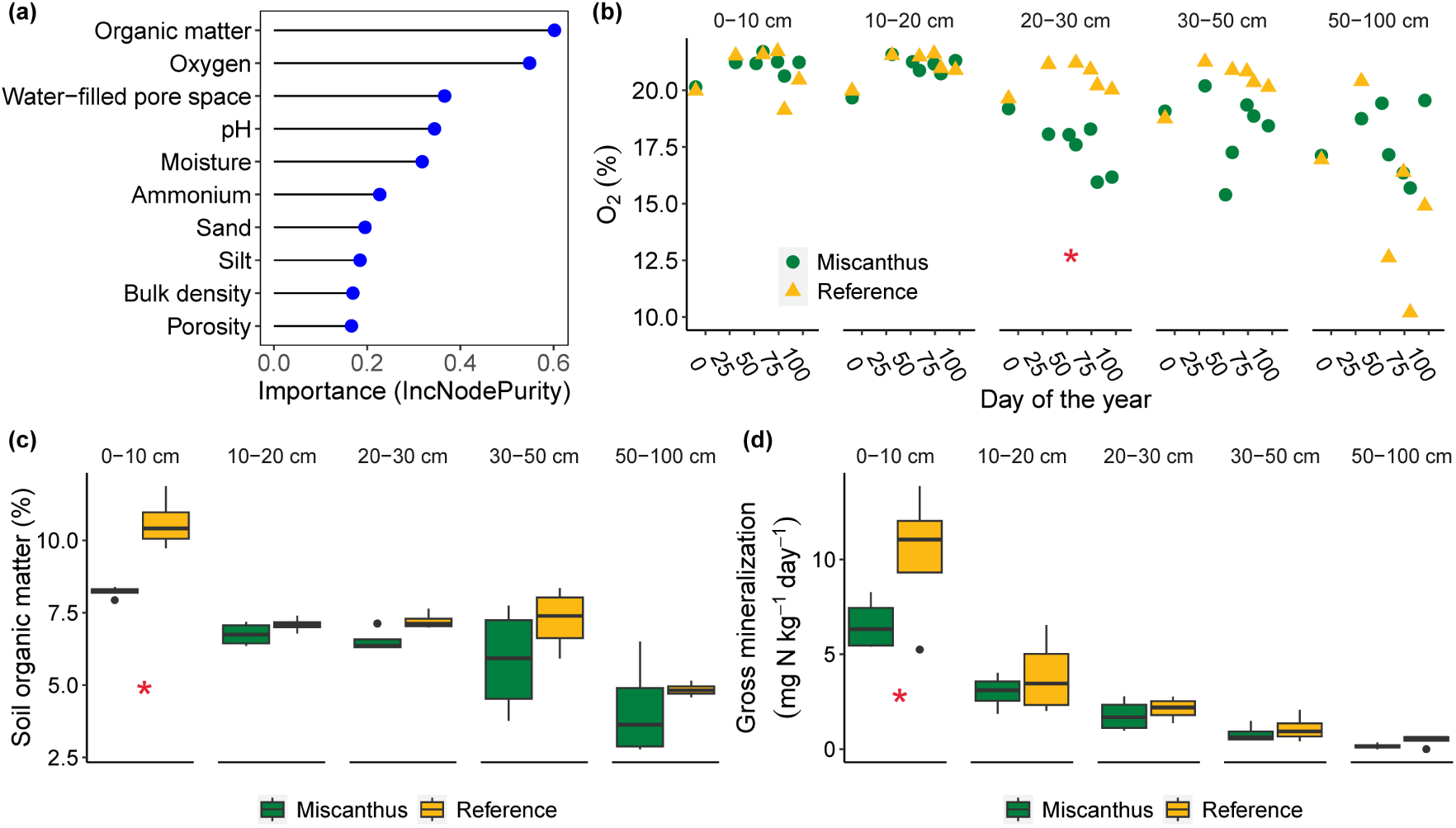
Factors shaping topsoil and subsoil AOA community. (a) The relative importance (increase in node purity) of soil properties in shaping AOA community differences between miscanthus and reference soil. (b) Soil O_2_ concentrations along miscanthus (green circles) and reference (yellow triangles) soil profiles. Each point represents a sampling date, which spans from December to April. (c, d) Soil organic matter content and gross N mineralization rates along miscanthus (green) and reference (yellow) soil profiles. Stars indicate significant differences between miscanthus and reference soil (*P < 0.05*). See Fig. 2 for boxplot interpretation.

Based on the biogeochemical function assigned by FAPROTAX, we found the functional abundance of aerobic heterotrophs was negatively correlated with the functional abundance of nitrification (constrained to AOA) for both miscanthus and reference subsoil (Fig. 4a, *P* < 0.05). However, much higher variation was explained in the reference than miscanthus subsoil (R^2^: 0.88 vs. 0.29). Moreover, the relationship between the aerobic heterotrophs’ functional abundance and the AOA’s functional abundance was significantly stronger in reference than miscanthus subsoil (Fig. 4a, slope: -1.98 vs. -0.80, *P* < 0.05). At the OTU level, DESeq2 analysis showed that miscanthus enriched more AOA in the subsoil than the shallow-rooted reference (*P < 0.01*) (Fig. 4b). AOA clades that were significantly different between miscanthus and reference soil belong to *Nitrososphaerales*-δ and *Nitrososphaerales*-γ. For aerobic heterotrophs, taxa that were depleted in the subsoil of miscanthus included the family *Sphingomonadaceae* and the genus *Solirubrobacter*, *Microvirga*, *Hyphomicrobium* and *Aeromicrobium* (*P < 0.01*, Fig. 4c). Aerobic heterotrophs that were enriched in miscanthus subsoil mostly belonged to the genus *Acidothermus* and *Nocardioides*.

**Fig. 4.**
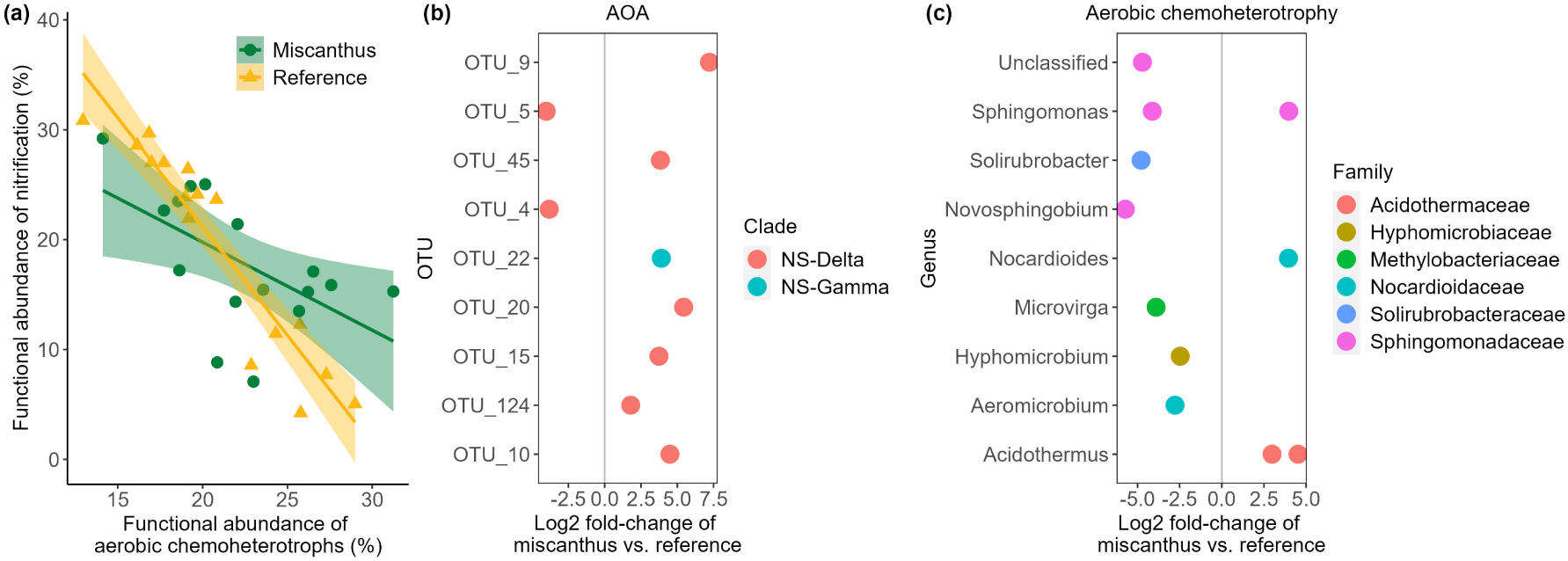
Microbial interactions in subsoil (10-100 cm depth). (a) Correlations between the functional abundance of aerobic heterotrophic bacteria and functional abundance of nitrification (constrained to AOA) in subsoil of miscanthus (green circle) and reference (yellow triangle). Shading represents 95% confidence interval. (b) Log2 fold change of AOA OTUs that are significantly enriched (right of vertical line) or depleted (left of vertical line) in the subsoil of miscanthus relative to reference (*P < 0.01*). Colors indicate clades of the taxonomic order *Nitrososphaerales* (NS). (c) Log2 fold change of aerobic heterotrophic bacteria (genus level) that are significantly enriched (right of vertical line) or depleted (left of vertical line) in the subsoil of miscanthus relative to reference (*P < 0.01*). Colors indicate taxonomic families.

## 4. Discussion

We investigated mechanisms regulating assembly and function of nitrifiers in topsoil versus subsoil by comparing soils underneath deep-rooted miscanthus versus adjacent shallow-rooted turfgrass references. In topsoil, lower SOM content and associated lower gross N mineralization resulted in strong tripartite competition among plants, heterotrophic bacteria and nitrifiers for available NH_4_^+^. This led to the suppression of community and function of AOA under miscanthus compared to the reference. In subsoil, suboxic conditions under miscanthus selected against aerobic heterotrophic bacteria, weakening the competition between AOA and heterotrophs to allow higher gross nitrification rates compared to under the turfgrass reference. Collectively, these findings suggest that plants can regulate archaeal nitrifier communities and function via altering SOM and O_2_ in topsoil and subsoil, respectively (Fig. 5).

**Fig. 5.**
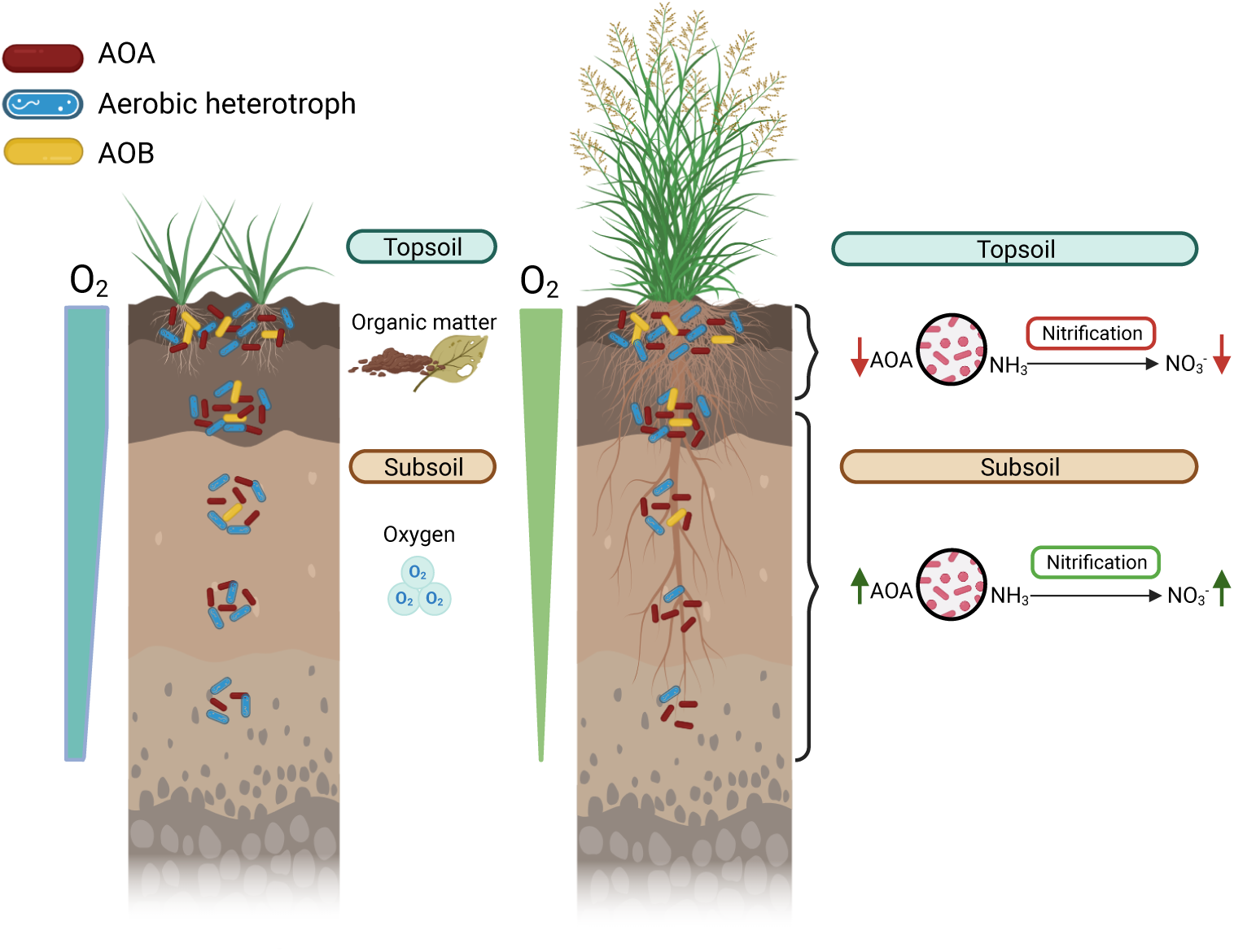
Conceptual diagram of microbial interactions along soil profiles under deep-rooted plants (right) versus shallow-rooted plants (left). In topsoil, the lower SOM and NH_4_^+^ supply under deep-rooted plants relative to shallow-rooted plants result in strong competition among plants, nitrifiers and heterotrophic bacteria for available N, leading to suppression of AOA and nitrification. Soil O_2_ decreases along soil profiles for both plant types, but suboxic conditions under deep-rooted plants, likely due to primed microbial activity and O_2_ consumption stimulated by rhizodeposition, favor O_2_-insensitive AOA over aerobic heterotrophic bacteria. As a result, nitrification is enhanced in subsoil of deep-rooted plants.

Soil organic matter was the most important factor determining AOA communities in the topsoil (Fig. 3a, 3c). Due to high root biomass near the soil surface (Sprunger et al., 2017), plant uptake of inorganic N in topsoil can be high (Xu et al., 2011), causing strong plant-microbe competition for the available N (Kuzyakov and Xu, 2013). With lower SOM content in topsoil, as observed for the miscanthus plots relative to the turfgrass alleyways at our study site (Fig. 3d), the decreased NH_4_^+^ supply from N mineralization (Fig. S7) can further intensify plant-heterotroph-nitrifier competition. Since nitrifiers are poor competitors for NH_4_^+^ compared to plant roots and heterotrophic bacteria (Zak et al., 1990; Verhagen et al., 1992), the intensified tripartite competition might have led to the suppressed nitrification of AOA in topsoil of miscanthus (Fig. 2). In comparison, once the nutrient limitation was relieved by the higher SOM and higher NH_4_^+^ supply, significantly higher gross nitrification and relative abundance of AOA were observed in reference topsoil (Fig. 2, Fig. S5).

In subsoil, deep-rooted miscanthus regulated AOA by altering soil O_2_ conditions (Fig. 3a, 3b). Compared to the reference, the significantly lower O_2_ from 20-50 cm depth in miscanthus may have stemmed from the primed microbial activity hence higher O_2_ consumption stimulated by rhizodeposition. As a result, significantly different AOA-aerobic heterotroph interactions between miscanthus and reference subsoil were observed (Fig. 4a). Compared to the steep negative correlations between functional abundance of AOA and functional abundance of aerobic chemoheterotrophs in reference subsoil (R^2^ = 0.88), a much weaker relationship was observed in miscanthus (R^2^ = 0.29). This difference could be explained by the differential tolerance of AOA vs. aerobic heterotrophs to soil O_2_. AOA has been suggested to require only 0.5 O_2_ per NH_3_ oxidized (Schleper and Nicol, 2010; Walker et al., 2010), and most of field surveys and culture-based studies have demonstrated that AOA could adapt to and function in O_2_-deficienct conditions (French et al., 2012; Pett-Ridge et al., 2013; Wang et al., 2014; Mushinski et al., 2017). Therefore, because low O_2_ conditions would favor AOA at the expense of weakened heterotrophic bacteria, a much weaker competition between AOA and aerobic heterotrophs was observed in miscanthus than reference subsoil (Fig. 4a). The weaker AOA-aerobic heterotroph competition under miscanthus also indicates AOA might have access to more available NH_4_^+^ in subsoil, which is consistent with the more enriched AOA (Fig. 4b) and higher gross nitrification rates in subsoil of miscanthus than reference (Fig. 2). In contrast, aerobic heterotrophs such as *Microvirga*, *Solirubrobacter* and *Aeromicrobium* were mostly depleted in miscanthus subsoil compared to reference (Fig. 4c), consistent with the preference of O_2_ by aerobic heterotrophs (Miller et al., 1991; Kanso and Patel, 2003; Singleton et al., 2003).

## 5. Conclusions

Overall, our study demonstrated that distinct mechanisms could drive plant-nitrifier interactions in topsoil and subsoil (Fig. 5). In topsoil, the strong plant-heterotroph-nitrifier competition induced by lower SOM and NH_4_^+^ supply could lead to the suppression of AOA. This relationship between SOM and nitrification has implications for predicting ecosystem services. While plants that accumulate SOM could confer climate change mitigation benefits, higher SOM can also increase nitrification rates and potentially increase soil NO_3_^-^ leaching and N_2_O emissions. In subsoil under deep-rooted plants, lower soil O_2_ can select against aerobic heterotrophs, liberating AOA from the strong heterotrophic competition for NH_4_^+^, resulting in higher nitrification rates compared to shallow-rooted plants. The role of O_2_ in mediating competition between AOA and heterotrophs for NH_4_^+^ is a newly identified mechanism by which deep-rooted plants can affect nitrification in subsoil. These findings demonstrate how plants can alter the soil environment differently in topsoil versus subsoil to mediate competitive interactions that affect nitrifier community structure and function.

## Acknowledgement

Funding for this work was provided by the DOE Center for Advanced Bioenergy and Bioproducts Innovation (U.S. Department of Energy, Office of Science, Office of Biological and Environmental Research under Award Number DE-SC0018420) and Future Interdisciplinary Research Explorations (FIRE) program of College of Agricultural, Consumer and Environmental Sciences (ACES), University of Illinois at Urbana-Champaign. D. Liang’s contribution was also supported by the Lawrence Livermore National Laboratory under the auspices of the U.S. DOE under Contract DE-AC52-07NA27344. We are grateful to A. Leakey and S. McCoy for the assistance with soil sampling and Giddings hydraulic system, and D. K. Lee for field site maintenance. We also thank R. V. Allen, H. Ware and J. Breiter for the field and lab assistance.

## Date availability

Sequence data used in this study are available in the NCBI Sequence Read Archive (accession number SUB12289181). All other data are contained in supplementary materials.

## References

Alves, R.J.E., Minh, B.Q., Urich, T., von Haeseler, A., Schleper, C., 2018. Unifying the global phylogeny and environmental distribution of ammonia-oxidising archaea based on *amoA* genes. Nature Communications 9, 1517. 10.1038/s41467-018-03861-1.

Apprill, A., McNally, S., Parsons, R., Weber, L., 2015. Minor revision to V4 region SSU rRNA 806R gene primer greatly increases detection of SAR11 bacterioplankton. Aquatic Microbial Ecology 75, 129–137. 10.3354/ame01753.

Banning, N.C., Maccarone, L.D., Fisk, L.M., Murphy, D.V., 2015. Ammonia-oxidising bacteria not archaea dominate nitrification activity in semi-arid agricultural soil. Scientific Reports 5, 11146. 10.1038/srep11146.

Bates, D., Mächler, M., Bolker, B., Walker, S., 2015. Fitting linear mixed-effects models using lme4. Journal of Statistical Software 67, 1–48. 10.18637/jss.v067.i01.

Bokulich, N.A., Kaehler, B.D., Rideout, J.R., Dillon, M., Bolyen, E., Knight, R., Huttley, G.A., Gregory Caporaso, J., 2018. Optimizing taxonomic classification of marker-gene amplicon sequences with QIIME 2’s q2-feature-classifier plugin. Microbiome 6, 90. 10.1186/s40168-018-0470-z.

Borcard, D., Gillet, F., Legendre, P., 2018. Association measures and matrices, In: Borcard, D., Gillet, F., Legendre, P. (Eds.), Numerical Ecology with R, Second ed. Springer New York, New York, NY, pp. 35-58.

Breiman, L., 2001. Random Forests. Machine Learning 45, 5–32. 10.1023/A:1010933404324.

Burnham, M.B., Simon, S.J., Lee, D., Kent, A.D., DeLucia, E.H., Yang, W.H., 2022. Intra- and inter-annual variability of nitrification in the rhizosphere of field-grown bioenergy sorghum. GCB Bioenergy 14, 393–410. 10.1111/gcbb.12917.

Button, E.S., Pett-Ridge, J., Murphy, D.V., Kuzyakov, Y., Chadwick, D.R., Jones, D.L., 2022. Deep-C storage: Biological, chemical and physical strategies to enhance carbon stocks in agricultural subsoils. Soil Biology and Biochemistry 170, 108697. 10.1016/j.soilbio.2022.108697.

Cooney, D.R., Namoi, N., Zumpf, C., Lim, S.-H., Villamil, M., Mitchell, R., Lee, D.K., 2022. Biomass production and nutrient removal by perennial energy grasses produced on a wet marginal land. Bioenergy Research. 10.1007/s12155-022-10488-0.

Daims, H., Lebedeva, E.V., Pjevac, P., Han, P., Herbold, C., Albertsen, M., Jehmlich, N., Palatinszky, M., Vierheilig, J., Bulaev, A., Kirkegaard, R.H., von Bergen, M., Rattei, T., Bendinger, B., Nielsen, P.H., Wagner, M., 2015. Complete nitrification by *Nitrospira* bacteria. Nature 528, 504–509. 10.1038/nature16461.

Diamond, S., Andeer, P.F., Li, Z., Crits-Christoph, A., Burstein, D., Anantharaman, K., Lane, K.R., Thomas, B.C., Pan, C., Northen, T.R., Banfield, J.F., 2019. Mediterranean grassland soil C–N compound turnover is dependent on rainfall and depth, and is mediated by genomically divergent microorganisms. Nature Microbiology 4, 1356–1367. 10.1038/s41564-019-0449-y.

Fish, J., Chai, B., Wang, Q., Sun, Y., Brown, C.T., Tiedje, J., Cole, J., 2013. FunGene: the functional gene pipeline and repository. Frontiers in Microbiology 4. 10.3389/fmicb.2013.00291.

French, E., Kozlowski Jessica, A., Mukherjee, M., Bullerjahn, G., Bollmann, A., 2012. Ecophysiological characterization of ammonia-oxidizing archaea and bacteria from freshwater. Applied and Environmental Microbiology 78, 5773–5780. 10.1128/AEM.00432-12.

Gee, G.W., Bauder, J.W., 1986. Particle-size Analysis, Methods of Soil Analysis, pp. 383–411.

Gill, R., Burke, I.C., Milchunas, D.G., Lauenroth, W.K., 1999. Relationship between root biomass and soil organic matter pools in the shortgrass steppe of Eastern Colorado. Ecosystems 2, 226–236. 10.1007/s100219900070.

Gillies, L.E., Thrash, J.C., deRada, S., Rabalais, N.N., Mason, O.U., 2015. Archaeal enrichment in the hypoxic zone in the northern Gulf of Mexico. Environmental Microbiology 17, 3847–3856. 10.1111/1462-2920.12853.

Hannonlab, 2010. FASTX-Toolkit: FASTQ/A short-reads pre-processing tools.

Hart, S.C., Stark, J.M., Davidson, E.A., Firestone, M.K., 1994. Nitrogen mineralization, immobilization, and nitrification, Methods of Soil Analysis, pp. 985–1018.

Hawkes, C.V., DeAngelis, K.M., Firestone, M.K., 2007. Root interactions with soil microbial communities and processes, In: Cardon, Z.G., Whitbeck, J.L. (Eds.), The Rhizosphere. Academic Press, Burlington, pp. 1–29.

Herman, D.J., Brooks, P.D., Ashraf, M., Azam, F., Mulvaney, R.L., 1995. Evaluation of methods for nitrogen-15 analysis of inorganic nitrogen in soil extracts. II. Diffusion methods. Communications in Soil Science and Plant Analysis 26, 1675–1685. 10.1080/00103629509369400.

Hoogsteen, M.J.J., Lantinga, E.A., Bakker, E.J., Groot, J.C.J., Tittonell, P.A., 2015. Estimating soil organic carbon through loss on ignition: effects of ignition conditions and structural water loss. European Journal of Soil Science 66, 320–328. 10.1111/ejss.12224.

Jones, D.L., Magthab, E.A., Gleeson, D.B., Hill, P.W., Sánchez-Rodríguez, A.R., Roberts, P., Ge, T., Murphy, D.V., 2018. Microbial competition for nitrogen and carbon is as intense in the subsoil as in the topsoil. Soil Biology and Biochemistry 117, 72–82. 10.1016/j.soilbio.2017.10.024.

Kanso, S., Patel, B.K.C., 2003. Microvirga subterranea gen. nov., sp. nov., a moderate thermophile from a deep subsurface Australian thermal aquifer. International Journal of Systematic and Evolutionary Microbiology 53, 401–406. 10.1099/ijs.0.02348-0.

Kaye, J.P., Hart, S.C., 1997. Competition for nitrogen between plants and soil microorganisms. Trends in Ecology & Evolution 12, 139–143. 10.1016/S0169-5347(97)01001-X.

Ke, X., Lu, W., Conrad, R., 2015. High oxygen concentration increases the abundance and activity of bacterial rather than archaeal nitrifiers in rice field soil. Microbial Ecology 70, 961–970. 10.1007/s00248-015-0633-4.

Krichels, A., DeLucia, E.H., Sanford, R., Chee-Sanford, J., Yang, W.H., 2019. Historical soil drainage mediates the response of soil greenhouse gas emissions to intense precipitation events. Biogeochemistry 142, 425–442. 10.1007/s10533-019-00544-x.

Kuzyakov, Y., Xu, X., 2013. Competition between roots and microorganisms for nitrogen: mechanisms and ecological relevance. New Phytologist 198, 656–669. 10.1111/nph.12235.

Lennon, J., 2020. Microbial life deep underfoot. mBio 11. 10.1128/mbio.03201-19.

Lenth, R.V., 2016. Least-squares means: the R package lsmeans. Journal of Statistical Software 69, 1–33. 10.18637/jss.v069.i01.

Liang, D., Ouyang, Y., Tiemann, L., Robertson, G.P., 2020. Niche differentiation of bacterial versus archaeal soil nitrifiers induced by ammonium inhibition along a management gradient. Frontiers in Microbiology 11. 10.3389/fmicb.2020.568588.

Louca, S., Parfrey, L.W., Doebeli, M., 2016. Decoupling function and taxonomy in the global ocean microbiome. Science 353, 1272–1277. 10.1126/science.aaf4507.

Love, M.I., Huber, W., Anders, S., 2014. Moderated estimation of fold change and dispersion for RNA-seq data with DESeq2. Genome Biology 15, 550. 10.1186/s13059-014-0550-8.

Miller, E.S., Woese, C.R., Brenner, S., 1991. Description of the erythromycin-producing bacterium *Arthrobacter* sp. strain NRRL B-3381 as *Aeromicrobium erythreum* gen. nov., sp. nov. International Journal of Systematic and Evolutionary Microbiology 41, 363–368. 10.1099/00207713-41-3-363.

Moreau, D., Bardgett, R.D., Finlay, R.D., Jones, D.L., Philippot, L., 2019. A plant perspective on nitrogen cycling in the rhizosphere. Functional Ecology 33, 540–552. 10.1111/1365-2435.13303.

Mushinski, R.M., Gentry, T.J., Dorosky, R.J., Boutton, T.W., 2017. Forest harvest intensity and soil depth alter inorganic nitrogen pool sizes and ammonia oxidizer community composition. Soil Biology and Biochemistry 112, 216–227. 10.1016/j.soilbio.2017.05.015.

Nogueira, R., Melo, L.s.F., Purkhold, U., Wuertz, S., Wagner, M., 2002. Nitrifying and heterotrophic population dynamics in biofilm reactors: effects of hydraulic retention time and the presence of organic carbon. Water Research 36, 469–481. 10.1016/S0043-1354(01)00229-9.

Oksanen, J., Simpson, G.L., Blanchet, F.G., Kindt, R., Legendre, P., Minchin, P.R., O’Hara, R.B., Solymos, P., Stevens, M.H.H., Szoecs, E., Wagner, H., Barbour, M., Bedward, M., Bolker, B., Borcard, D., Carvalho, G., Chirico, M., Caceres, M.D., Durand, S., Evangelista, H.B.A., FitzJohn, R., Friendly, M., Furneaux, B., Hannigan, G., Hill, M.O., Lahti, L., McGlinn, D., Ouellette, M.-H., Cunha, E.R., Smith, T., Stier, A., Braak, C.J.F.T., Weedon, J., 2022. vegan: community ecology package.

Ouyang, Y., Norton, J.M., Stark, J.M., Reeve, J.R., Habteselassie, M.Y., 2016. Ammonia-oxidizing bacteria are more responsive than archaea to nitrogen source in an agricultural soil. Soil Biology and Biochemistry 96, 4–15. 10.1016/j.soilbio.2016.01.012.

Parada, A.E., Needham, D.M., Fuhrman, J.A., 2016. Every base matters: assessing small subunit rRNA primers for marine microbiomes with mock communities, time series and global field samples. Environmental Microbiology 18, 1403–1414. 10.1111/1462-2920.13023.

Pertea, M., Robeson, M.S., O’Rourke, D.R., Kaehler, B.D., Ziemski, M., Dillon, M.R., Foster, J.T., Bokulich, N.A., 2021. RESCRIPt: Reproducible sequence taxonomy reference database management. PLoS Computational Biology 17. 10.1371/journal.pcbi.1009581.

Pett-Ridge, J., Petersen, D.G., Nuccio, E., Firestone, M.K., 2013. Influence of oxic/anoxic fluctuations on ammonia oxidizers and nitrification potential in a wet tropical soil. FEMS Microbiology Ecology 85, 179–194. 10.1111/1574-6941.12111.

Philippot, L., Hallin, S., Börjesson, G., Baggs, E.M., 2009. Biochemical cycling in the rhizosphere having an impact on global change. Plant and Soil 321, 61–81. 10.1007/s11104-008-9796-9.

Prosser, J.I., 1990. Autotrophic nitrification in bacteria, In: Rose, A.H., Tempest, D.W. (Eds.), Advances in Microbial Physiology. Academic Press, pp. 125–181.

Prosser, J.I., Hink, L., Gubry-Rangin, C., Nicol, G.W., 2019. Nitrous oxide production by ammonia oxidizers: Physiological diversity, niche differentiation and potential mitigation strategies. Global Change Biology 26, 103–118. 10.1111/gcb.14877.

Prosser, J.I., Nicol, G.W., 2012. Archaeal and bacterial ammonia-oxidisers in soil: the quest for niche specialisation and differentiation. Trends in Microbiology 20, 523–531. 10.1016/j.tim.2012.08.001.

Quast, C., Pruesse, E., Yilmaz, P., Gerken, J., Schweer, T., Yarza, P., Peplies, J., Glöckner, F.O., 2013. The SILVA ribosomal RNA gene database project: improved data processing and web-based tools. Nucleic Acids Research 41, D590–D596. 10.1093/nar/gks1219.

Robertson, G.P., Groffman, P.M., 2015. Nitrogen transformations, In: Paul, E.A. (Ed.), Soil Microbiology, Ecology and Biochemistry, Fourth ed. Academic Press, Burlington, pp. 421–446.

Rognes, T., Flouri, T., Nichols, B., Quince, C., Mahé, F., 2016. VSEARCH: a versatile open source tool for metagenomics. PeerJ 4. 10.7717/peerj.2584.

Rotthauwe, J.H., Witzel, K.P., Liesack, W., 1997. The ammonia monooxygenase structural gene amoA as a functional marker: molecular fine-scale analysis of natural ammonia-oxidizing populations. Applied and Environmental Microbiology 63, 4704–4712. 10.1128/aem.63.12.4704-4712.1997.

Salomé, C., Nunan, N., Pouteau, V., Lerch, T.Z., Chenu, C., 2010. Carbon dynamics in topsoil and in subsoil may be controlled by different regulatory mechanisms. Global Change Biology 16, 416–426. 10.1111/j.1365-2486.2009.01884.x.

Schleper, C., Nicol, G.W., 2010. Ammonia-oxidising archaea – physiology, ecology and evolution, In: Poole, R.K. (Ed.), Advances in Microbial Physiology. Academic Press, pp. 1–41.

Silver, W.L., Lugo, A.E., Keller, M., 1999. Soil oxygen availability and biogeochemistry along rainfall and topographic gradients in upland wet tropical forest soils. Biogeochemistry 44, 301–328. 10.1007/BF00996995.

Singleton, D.R., Furlong, M.A., Peacock, A.D., White, D.C., Coleman, D.C., Whitman, W.B., 2003. *Solirubrobacter pauli* gen. nov., sp. nov., a mesophilic bacterium within the *Rubrobacteridae* related to common soil clones. International Journal of Systematic and Evolutionary Microbiology 53, 485–490. 10.1099/ijs.0.02438-0.

Sprunger, C.D., Oates, L.G., Jackson, R.D., Robertson, G.P., 2017. Plant community composition influences fine root production and biomass allocation in perennial bioenergy cropping systems of the upper Midwest, USA. Biomass and Bioenergy 105, 248–258. 10.1016/j.biombioe.2017.07.007.

Subbarao, G.V., Nakahara, K., Hurtado, M.P., Ono, H., Moreta, D.E., Salcedo, A.F., Yoshihashi, A.T., Ishikawa, T., Ishitani, M., Ohnishi-Kameyama, M., Yoshida, M., Rondon, M., Rao, I.M., Lascano, C.E., Berry, W.L., Ito, O., 2009. Evidence for biological nitrification inhibition in Brachiaria pastures. Proceedings of the National Academy of Sciences 106, 17302–17307. 10.1073/pnas.0903694106.

Tess, B., Aronson Emma, L., Arogyaswamy, K., Billings Sharon, A., Botthoff Jon, K., Campbell Ashley, N., Dove Nicholas, C., Fairbanks, D., Gallery Rachel, E., Hart Stephen, C., Kaye, J., King, G., Logan, G., Lohse Kathleen, A., Maltz Mia, R., Mayorga, E., O’Neill, C., Owens Sarah, M., Packman, A., Pett-Ridge, J., Plante Alain, F., Richter Daniel, D., Silver Whendee, L., Yang Wendy, H., Fierer, N., 2019. Ecological and genomic attributes of novel bacterial taxa that thrive in subsurface soil horizons. mBio 10, e01318–01319. 10.1128/mBio.01318-19.

Tilman, G.D., 1984. Plant dominance along an experimental nutrient gradient. Ecology 65, 1445–1453. 10.2307/1939125.

Tourna, M., Freitag, T.E., Nicol, G.W., Prosser, J.I., 2008. Growth, activity and temperature responses of ammonia-oxidizing archaea and bacteria in soil microcosms. Environmental Microbiology 10, 1357–1364. 10.1111/j.1462-2920.2007.01563.x.

Trivedi, C., Reich, P.B., Maestre, F.T., Hu, H.-W., Singh, B.K., Delgado-Baquerizo, M., 2019. Plant-driven niche differentiation of ammonia-oxidizing bacteria and archaea in global drylands. The ISME Journal 13, 2727–2736. 10.1038/s41396-019-0465-1.

Van den Berge, K., Perraudeau, F., Soneson, C., Love, M.I., Risso, D., Vert, J.-P., Robinson, M.D., Dudoit, S., Clement, L., 2018. Observation weights unlock bulk RNA-seq tools for zero inflation and single-cell applications. Genome Biology 19, 24. 10.1186/s13059-018-1406-4.

van Kessel, M.A.H.J., Speth, D.R., Albertsen, M., Nielsen, P.H., Op den Camp, H.J.M., Kartal, B., Jetten, M.S.M., Lücker, S., 2015. Complete nitrification by a single microorganism. Nature 528, 555–559. 10.1038/nature16459.

Verhagen, F., Duyts, H., Laanbroek Hendrikus, J., 1992. Competition for ammonium between nitrifying and heterotrophic bacteria in continuously percolated soil columns. Applied and Environmental Microbiology 58, 3303–3311. 10.1128/aem.58.10.3303-3311.1992.

Walker, C.B., de la Torre, J.R., Klotz, M.G., Urakawa, H., Pinel, N., Arp, D.J., Brochier-Armanet, C., Chain, P.S.G., Chan, P.P., Gollabgir, A., Hemp, J., Hügler, M., Karr, E.A., Könneke, M., Shin, M., Lawton, T.J., Lowe, T., Martens-Habbena, W., Sayavedra-Soto, L.A., Lang, D., Sievert, S.M., Rosenzweig, A.C., Manning, G., Stahl, D.A., 2010. *Nitrosopumilus maritimus* genome reveals unique mechanisms for nitrification and autotrophy in globally distributed marine crenarchaea. Proceedings of the National Academy of Sciences 107, 8818–8823. 10.1073/pnas.0913533107.

Wang, B., Zhao, J., Guo, Z., Ma, J., Xu, H., Jia, Z., 2015. Differential contributions of ammonia oxidizers and nitrite oxidizers to nitrification in four paddy soils. The ISME Journal 9, 1062–1075. 10.1038/ismej.2014.194.

Wang, J., Zhang, L., Lu, Q., Raza, W., Huang, Q., Shen, Q., 2014. Ammonia oxidizer abundance in paddy soil profile with different fertilizer regimes. Applied Soil Ecology 84, 38–44. 10.1016/j.apsoil.2014.06.009.

Xu, X., Ouyang, H., Richter, A., Wanek, W., Cao, G., Kuzyakov, Y., 2011. Spatio-temporal variations determine plant–microbe competition for inorganic nitrogen in an alpine meadow. Journal of Ecology 99, 563–571. 10.1111/j.1365-2745.2010.01789.x.

Zak, D.R., Groffman, P.M., Pregitzer, K.S., Christensen, S., Tiedje, J.M., 1990. The vernal dam: plant-microbe competition for nitrogen in northern hardwood forests. Ecology 71, 651–656. 10.2307/1940319.

Zumpf, C., Lee, M.S., Thapa, S., Guo, J., Mitchell, R., Volenec, J.J., Lee, D., 2019. Impact of warm-season grass management on feedstock production on marginal farmland in Central Illinois. GCB Bioenergy 11, 1202–1214. 10.1111/gcbb.12627.

